# Human Biopsies in Nanofibrillar Cellulose Hydrogel – A Novel Method for Long-term Tissue Culture

**DOI:** 10.1101/2021.11.22.466872

**Authors:** Johanna Niklander, Raili Koivuniemi, Alexander Stallinger, Florian Kleinegger, Lauri Paasonen, Silke Schrom, Bernadette Liegl-Atzwanger, Iris Zalaudek, Gord von Campe, Georg Singer, Johannes Haybaeck, Marjo Yliperttula, Beate Rinner

## Abstract

Advanced 3D in vitro models are laborious to prepare and susceptible to unintentional design errors due to culture adaptations, cell immaturity, xenofactors or yet incomplete knowledge of the dynamics within tissues or materials. In order to acquire cost-efficient research material with intact in vivo composition, we developed novel tissue culture method with plant-derived scaffolding.

Human skin-, foreskin- and glioblastoma multiforme biopsies were dissected mechanically and cultivated for 28 days in plant-derived nanofibrillar cellulose hydrogel. Comparative cultures were done using mouse sarcoma tumor –derived Matrigel™. Long-term preservation of cultivated tissues was evaluated against typical immunohistochemical biomarkers for each tissue type: skin tissues for cytokeratins 5/6, E-cadherin and vimentin for sustained tissue structures, and brain neoplasia for Olig2, S100, Nestin, NOTCH1, MAP2 and GFAP for preserved disease profile.

Histological analysis from both culture conditions showed that until day 28, all cultivated biopsy types were able to sustain their characteristic protein expressions without signs of necrosis. We here conclude a novel tissue culture model in xeno-free 3D scaffolding, that can enable long-term sample storage in vitro, studies of human tumor tissues and their non-neoplastic microenvironment, and innovations in personalized medicine research.

## Introduction

Most of the current cell models consist of one or two cell types, whereas to be physiologically relevant, the model should also take into account the complex cellular interactions within tissue microenvironment [1, 2]. However, highly complex in vitro models are laborious [3, 4], and may accumulate unintentional design errors [5]. Culture materials used enabling 3D growth have also been known to affect the diffusion environment of experiments [6], introducing xenofactor contaminations to results [7, 8] and affecting cell behavior via scaffold stiffness or growth factors [9, 10]. Formation of physiological structures that require growth in cell aggregate size also face limitations due to limited nutrient and oxygen diffusion in 3D cell cultures [11, 12, 13].

Due to many challenges in reconstructing disease models in vitro, ex vivo samples have been gaining solid ground as experimental material. Intact biopsy pieces are easy and fast to process for cultivation, readily contain all cell populations of the studied biological target and have the genetic makeups of patients. Tissue in vitro applications have been successfully utilized in Dichelobacter nodosus pathogenesis modelling using ovine skin cultures and agarose scaffolding [14], in the evaluation of novel photodynamic therapy with human skin biopsy cultures [15], and in the evaluation of human tumor metabolism responses to experimental drug in alternative oxygen tensions [16]. Further, patient-derived material has brought clinically relevant improvements in personalized medicine research and pharmacodynamics modelling [17]. When reliably validated for general use, the robustness of setting up a 3D biopsy culture model could significantly save time, costs and resources in the industry.

Greatest challenge in tissue sample culture is to sustain all relevant cell types long-term in culture conditions [11]. In this study, our aim was to establish long-term tissue cultures from human biopsy pieces, and to evaluate the preservation of tissue-typical protein expressions. Experimental human biopsy samples were acquired from healthy skin (HS)-, healthy foreskin (FRSK)- and glioblastoma multiforme (GBM) tissue donors. In order to avoid unintentional regulatory contacts as well as to qualify experimental culture model for future clinical prospects, we utilized nanofibrillar cellulose hydrogel as culture scaffold (NFC; GrowDex®) [18, 19, 20]. Wood-derived, biocompatible NFC is composed of cellulose fibrils having nanoscale diameter and microscale length, and the fibril size and viscoelastic properties resemble those of extracellular matrix (ECM) [21]. Experiments with labeled dextrans have indicated NFC as a similar diffusion environment to human ECM [21]. To observe how NFC compares to a commonly known animal-derived culture scaffold, parallel biopsy cultures were done with Engelbreth-Holm-Swarm (EHS) mouse sarcoma ECM gel (BD Matrigel™) [10, 17]. Matrigel consists of laminin-111 (60%), collagen IV (30%), entactin (8%), perlecan and various growth factors [22,23], and provides cells a growth niche reminiscent to the natural basal lamina layer of ECM [24].

Patient biopsies were dissected into cultivated pieces using surgical scissors and inserted to NFC and Matrigel scaffoldings. Morphological development of the cultures was monitored with light microscopy, and histological correspondence with original ex vivo biopsy compositions was examined with immunohistochemical (IHC) profiling. Normal skin and foreskin tissues were evaluated for cytokeratins 5 and 6 (CK 5/6), epidermal E-cadherin, and mesenchymal vimentin in order to evaluate tissue structure survival. Glioblastoma samples were evaluated for typical histopathological disease profile using Olig2, S100, Nestin, NOTCH1, MAP2 and GFAP biomarkers.

Here, we present that human skin- and brain tissue samples sustained their histological profiles for 28 days when cultivated in NFC- and Matrigel matrices, and that these culture methods enabled constant morphology and histology characterization. The suitability of NFC in tissue culture enables reducing the use of species-different substances in experimental settings.

## Materials and Methods

### Patient biopsies

Ethical approval was in accordance with the ethical standards of the responsible committee for human experimentation and the current version of the Declaration of Helsinki. Ethical permissions 28-476ex15/16 and 26-219ex13/14 were used to obtain a HS biopsy, juvenile FRSK-1, −2 -and 3 biopsies via circumcision from patients younger than six years, and a GBM biopsy. Anonymized patient samples were received directly after surgery in 30 ml DMEM solution that contained minor blood and tissue fluid remnants. Biopsies were processed for 3D cultures within 3h post-operation, and were stored at +4°C before processing. Each patient was considered as an experimental repeat, and each cultivated tissue sample derived from the same original biopsy piece, a biological replicate (presented experimental repeats for NFC: HS n=1, FRSK n=3, GBM n=1; Matrigel: HS n=1, FRSK n=1, GBM n=1).

### Preparation of NFC- and Matrigel 3D culture scaffolds

Culture scaffold components were combined according to Table 1, where culture medium consisted of 10% (v/v) FBS (Biochrom AG, Berlin, Germany), 88% (v/v) RPMI-1640-31870 (Gibco, USA), 1mM L-glutamine (Life Technologies) and 1x P/S Pen/Strep (Life Technologies). Low retention pipette tips and utilization of tip graduations were found recommendable for NFC and Matrigel pipetting. Due to Matrigel’s instability at room temperature (RT), pipette tips and tube racks were cooled in freezer temperature before use, and culture scaffolds were prepared as individual workflows.

**Table 1.** Nanofibrillar Cellulose Hydrogel (NFC; GrowDex) and Matrigel culture scaffold components.

| GrowDex<br>(UPM Biomedicals,<br>Finland) | GrowDex<br>stock<br>concentration | Final GrowDex<br>concentration | GrowDe<br>x<br>volume | Culture<br>medium<br>volume | Ready<br>GrowDex<br>scaffold<br>volume |
| --- | --- | --- | --- | --- | --- |
|  | 1.5% | 0.7% (All cultures) | 500 µl | 571 µl | 1071 µl |
| Matrigel<br>Growth Factor<br>Reduced (GFR)<br>(BD Biosciences,<br>U.S.) | Matrigel stock<br>concentration<br>(protein mg) | Final Matrigel<br>concentration (v/v) | Matrigel<br>volume |  | Ready<br>Matrigel<br>scaffold<br>volume |
|  | 9.6 mg / ml | ≈40% (GBM*) | 500 µl | 750 µl | 1250 µl |
|  | 9.6 mg / ml | ≈70% (HS**, FRSK-1,2,3***) | 500 µl | 214 µl | 714 µl |

\*Glioblastoma multiforme biopsy, \*\*Healthy skin biopsy, \*\*\*Healthy foreskin biopsy 1, -2, -and 3.

Pre-calculated stock volumes of NFC and ice-cold Matrigel were pipetted into 2 ml sterile Eppendorf tubes, where the tube receiving Matrigel was holding pre-calculated volume of ice-cold culture medium and was positioned in a frozen tube rack or in ice. The pre-calculated medium volume for the NFC scaffold was pipetted directly into the hydrogel after the addition of NFC stock volume to tube. Both scaffolds were mixed with careful pipetting within the Eppendorf tubes (minimum of 90 seconds for the NFC), avoiding raising the materials high within the pipette tips. Possible air bubbles could be removed by tapping the tubes or by careful vortexing (NFC), and scaffolds were considered homogenous when no visible scaffold stock matrices could be detected. Ready NFC scaffold could be stored in RT, or within sterile 50 ml tube at +37°C, if used within same day, and at +4°C for one week, if no unstable culture medium components were added. Remaining Matrigel scaffold mixture was stored at +4°C on ice, and was used within the same day.

### 3D culture assembly

Preceding biopsy insertions, 50 μl of ready NFC- and Matrigel culture scaffolds were added to 96-culture well bottoms and placed into a cell incubator for a minimum of 30 min (NFC) and 1h (Matrigel). Before inserted to culture scaffolds, a biopsy was transferred to 10% v/v amphotericin B - Pen-Strep – PBS(-) solution for 5-7 min RT incubation, from which after the biopsy was rinsed in PBS(-) and moved into a 6-well plate holding enough warm DMEM to cover the tissue. Using sterile surgical scissors and tweezers, the biopsy was dissected into an average of 1.5-2 mm pieces that were inserted as flat tissue layers on top of the pre-incubated NFC- and Matrigel scaffold layers. Additional 50 μl of NFC- or Matrigel culture scaffolds were added on top of the inserted biopsy pieces, and the assembled 3D culture plates were further incubated for a minimum of 30 min (NFC) and 45 min (Matrigel) in a cell incubator. After incubations, 130 μl of culture medium was carefully added through a well wall on top of each culture well scaffold, where tilting of the NFC culture plate was found recommendable in order to minimize flow disturbance to scaffold layers. Suspension cultures were also done to acquire visual control condition for tissue viability after the ap-plied antibiotic treatment, and to observe cell outgrowth rate and variation in cell populations with the used medium exchange intervals. Additional liquid reservoir wells were found to be recommendable to avoid medium evaporation. Experimental setup is illustrated in Fig 1A,B.

**Fig 1.**
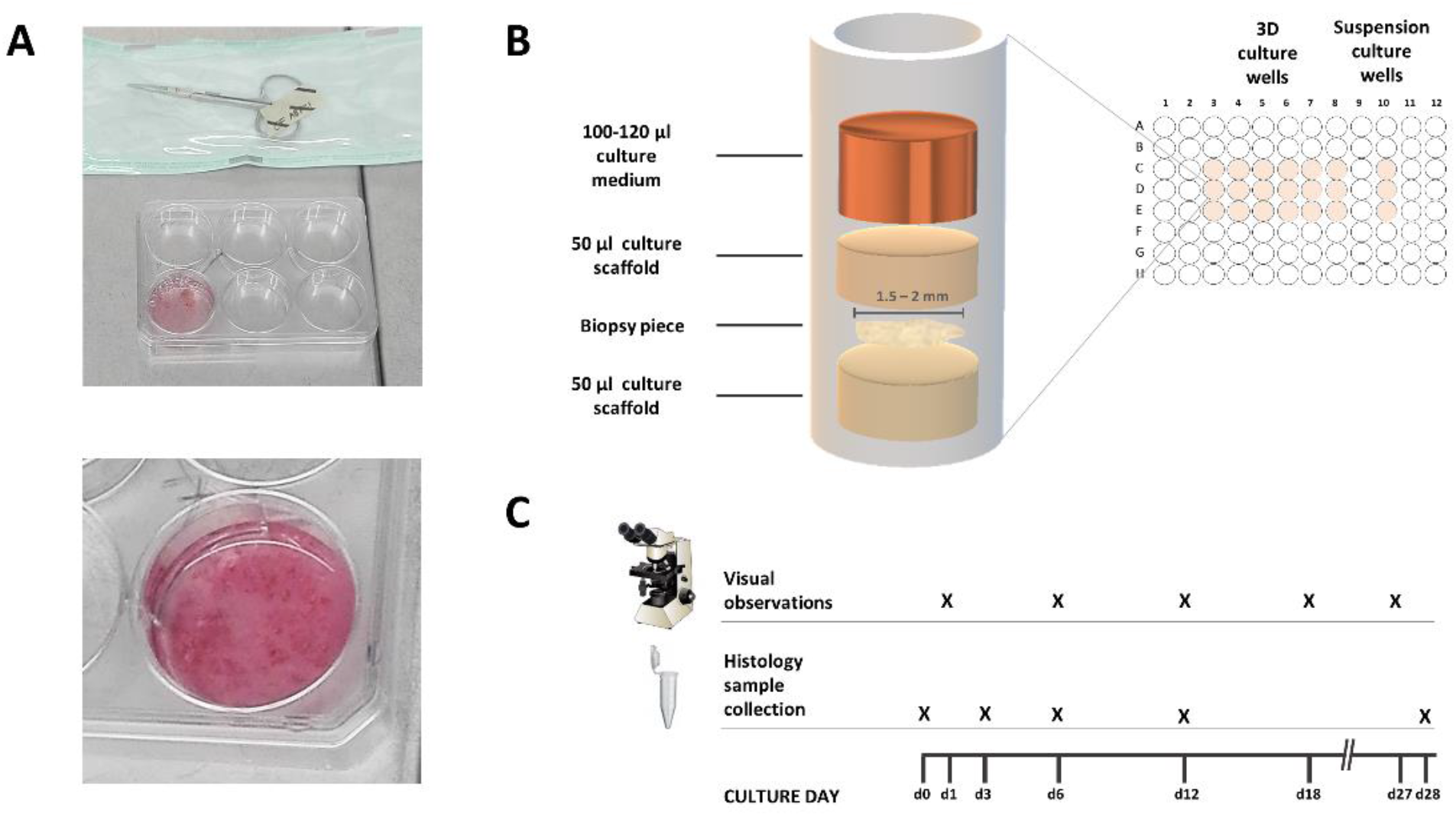
Experimental set up. Human healthy skin-, foreskin- and glioblastoma multiforme biopsies were treated with antibiotics and dissected in +37°C medium using sterile scissors and tweezers (A). Dissected biopsy pieces were implanted as flat tissue layers into 96-culture wells holding 50 μl layers of pre-incubated (+37°C) nanofibrillar cellulose (NFC) - or Matrigel culture scaffolding and were covered by another 50 μl layer of the same scaffolding. The assembled 3D culture plates were incubated once more in +37°C, from which after 130 μl of culture medium was added on top of the uppermost scaffold layers and was refreshed with 100-120 μl culture medium volume every three days. Suspension cultures were used as visual controls to observe sample viability, cell type variation and cell outgrowth with the applied antibiotic treatment and medium exchange intervals (B). Culture morphology development was observed with a light microscope on days 1, 6, 12, 18 and 27. Biopsy pieces were collected for histological analysis during ex vivo biopsy dissection (d0) and from 3D culture wells on days 3, 6, 12 and 28 (C).

### 3D culture maintenance

Culture medium was changed every 3 days from the topmost medium layer, where spent medium was refreshed to sustain ca. 100-120 μl medium volume during cultivation (Fig 1B). The first medium change was done on day 3, and repeated on the average every three days onwards in respect to culture wells’ pH indicator (phenol red). Morphology development of cultivated biopsy pieces was observed with a light microscope (Olympus TH4-200, Olympus America Inc.; USA) on days 1, 6, 12, 18 and 27 using cellSens imaging software (Olympus, USA) (Fig 1C).

### Histology sample collection

Biopsy samples were collected on days 0, 3, 6, 12 and 28 (Fig 1C). Upon 3D culture assembly, dissected ex vivo biopsy pieces were briefly rinsed in PBS(-) and incubated in 4% formalin at RT overnight, and then at 4°C for 24h. Cultivated GBM and FRSK-1 samples were collected from NFC by degrading the surrounding scaf-folding with an overnight GrowDase^®^ incubation (UPM Biomedicals, Finland), where used enzyme concentration was 140-200 mg/g (w/w) of GrowDase for each individual culture well scaffold. On the following day, GrowDase-treated tissue samples were col-lected with a pipette tip for 2x 3 min PBS(-) rinses, and treated with formalin incubations similarly to the ex vivo samples. During methodology development, GrowDase treatment was found unnecessary for collecting NFC-cultivated skin tissues, and was discontinued before HS- and FRSK-2 and −3 cultures. Without the use of GrowDase, NFC- and Matrigel –cultivated samples were collected directly from the culture well scaffolds with a pipette tip, followed by brief PBS(-) rinse and the described formalin incubations. Fixed samples were stored in 70% EtOH at 4°C before paraffinization. In order to prevent sample losses during automated paraffin processing, formalin-preserved pieces were rinsed from EtOH remnants and encapsulated in Richard-Allan Scientific™ HistoGel (Thermo Fisher, US). Liquidized HistoGel was solidified into a parafilmlined Eppendorf lid as a 30 μl layer, from which after a pre-fixed and rinsed biopsy piece was placed on top of the gel. Another 30 μl layer of skin-warm HistoGel was applied on top of the biopsy piece, and solidified at RT. HistoGel-embedded samples were further incubated in 4% formalin for 2h before transferring to paraffin-embedding sample cassettes and paraffin processing equipment VIP Tissue Tek 5E (Sakura, Japan).

### Immunohistochemistry

Histological analyses were done in the Diagnostic and Research Institute of Pathology, Medical University of Graz. The ex vivo- and cultivated biopsy samples were stained with hematoxylin-eosin (HE) to investigate the present tissue structures in each sample, and evaluated with antibodies characteristic for each tissue type (Table 2).

**Table 2.**
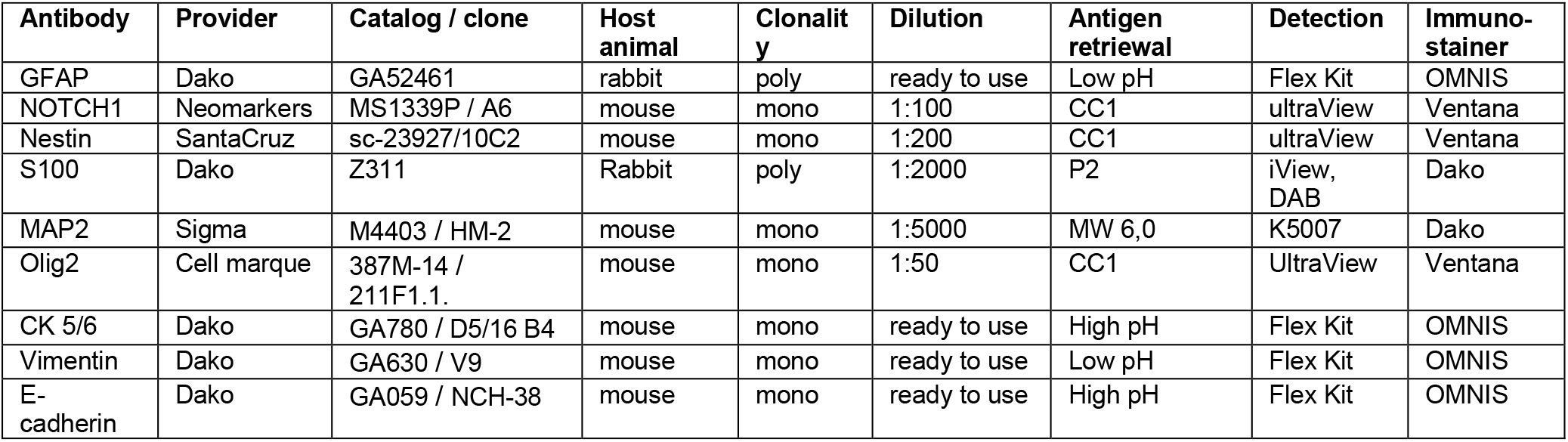
Antibodies used in immunohistochemistry analysis.

## Results

### Visual observations of HS and FRSK skin tissues revealed differing behaviors in vitro

On a general level, HS biopsies appeared relatively static during NFC cultivation, and development in the biopsy morphology was observable in a single well (Fig 2A-B). Visual observations were found to be limited in NFC cultures due to material’s opaque nature (Fig 2I), comparing to Matrigel culture wells (Fig 2J). During Matrigel cultivation, early rearrangements in the biopsy shape (Fig 2: E-H) and modest cell spreading could be observed (Fig 2: F-H, K). Occasional Matrigel-implanted HS biopsies were surrounded by transparent formations that contained particles resembling migrating cells (Fig 2J).

**Fig 2.**
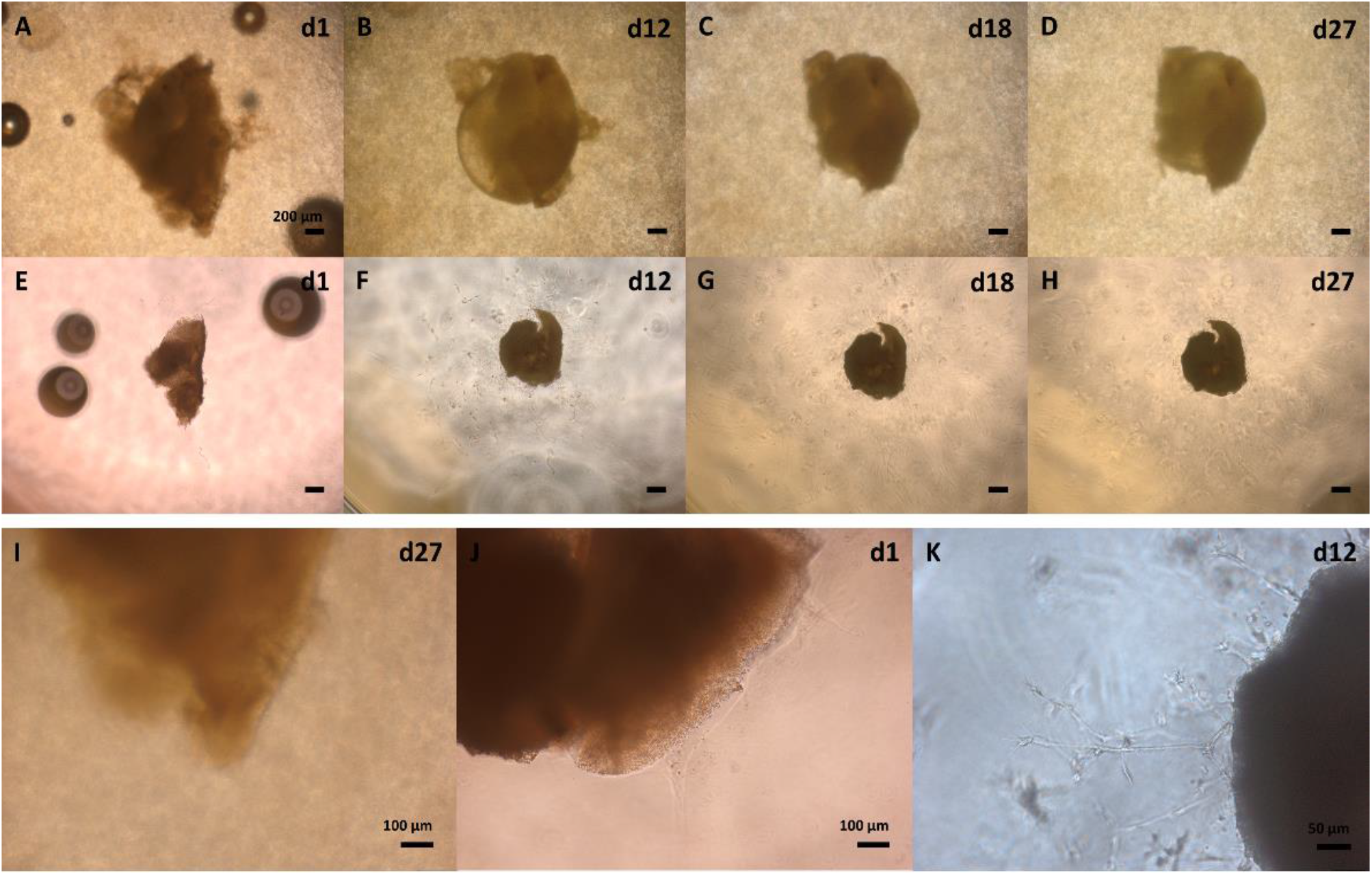
Morphology development of cultivated healthy skin (HS) biopsy pieces in 0.7% nanofibrillar cellulose (NFC) and 70% Matrigel. HS tissue showed minor size reductions during NFC cultivation (A-D), and new growth consisting of transparent tissue mass was observed on day 12 (B). Matrigel-cultivated tissues demonstrated permanent shape reformations in early culture stages and minor to none size reductions during cultivation (E-H). Modest cell spreading was commonly observed without noticeable size reductions to biopsy pieces (K). Opaque nature of NFC caused challenges to the visual observations of fine structures (I). Visual observations were clear in Matrigel and transparent rims containing cell-like particles could be detected surrounding the Matrigel-implanted biopsy pieces (J). Figure scale bars 200 μm (A-H) 100 μm (I-J), 50 μm (K).

NFC-cultivated FRSK tissues showed growth of tissue mass, development in the biopsy shape (Fig 3A-D), and signs of protruding or degrading structures (Fig 3C-D,I). When cultivated in Matrigel scaffolding, rapid cell spreading, tissue growth and biopsy unfolding were observed (Fig 3J,E-H). Reductions in biopsy size correlated with rapid cell spreading in the Matrigel cultures. No noticeable amounts of dying cells or other debris were detected with either skin tissue types after the first three days of culture in NFC or Matrigel.

**Fig 3.**
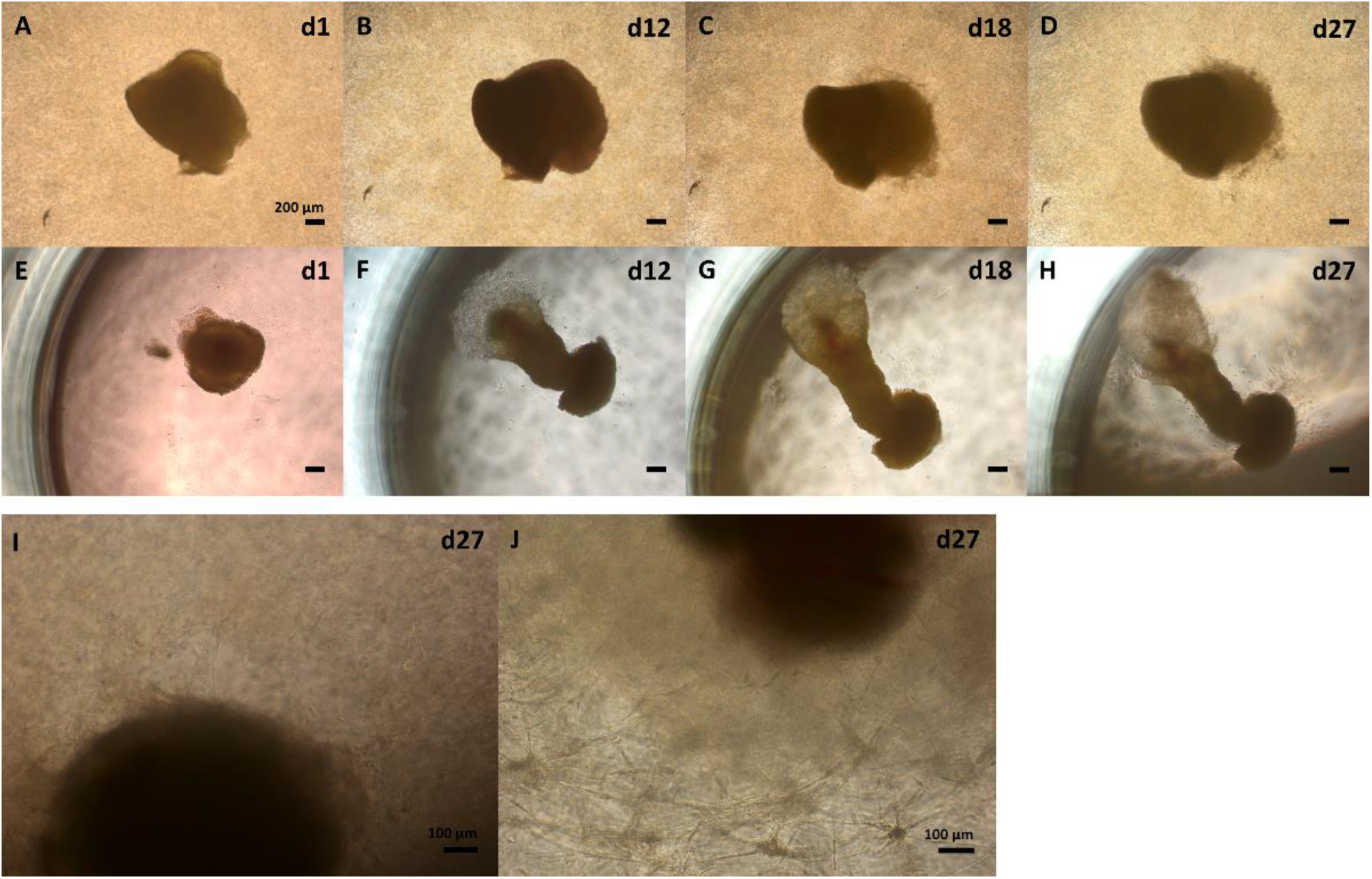
Morphology development of cultivated human foreskin (FRSK) biopsy pieces in 0.7% nanofibrillar cellulose (NFC) and 70% Matrigel. Growth of new tissue mass and development in biopsy pieces’ shape were observed during NFC culture (A-D). Undefined structures resembling spreading growth or tissue degradation were also observed in the NFC cultures (C, D, I). Occasional culture wells demonstrated biopsy unfolding and tissue growth during Matrigel cultivation (E-H). Protrusive tissue structures in surrounding culture scaffold were rarely observed in opaque NFC (I), and frequently in clear Matrigel scaffolding (J). Figure scale bars 200 μm (A-H), 100 μm (I-J).

### Visual observations of GBM biopsies showed cell outgrowth

GBM cultures showed greatest variation in the growth rates of individual culture wells and most prominent size reductions in cultivated tissue pieces. Typical morphology development during both NFC and Matrigel cultivation included partial degradation of original biopsy structures (Fig 4A-D) and appearance of protruding cells (Fig 4I,J,K). Extensive cell spreading in 3D Matrigel correlated with biopsy size reductions (Fig 4E-H). Dying cells caused low visibility to many culture wells between days 6-12. The turbid medium was clearing spontaneously during cultivation without additional medium changes or antibiotics.

**Fig 4.**
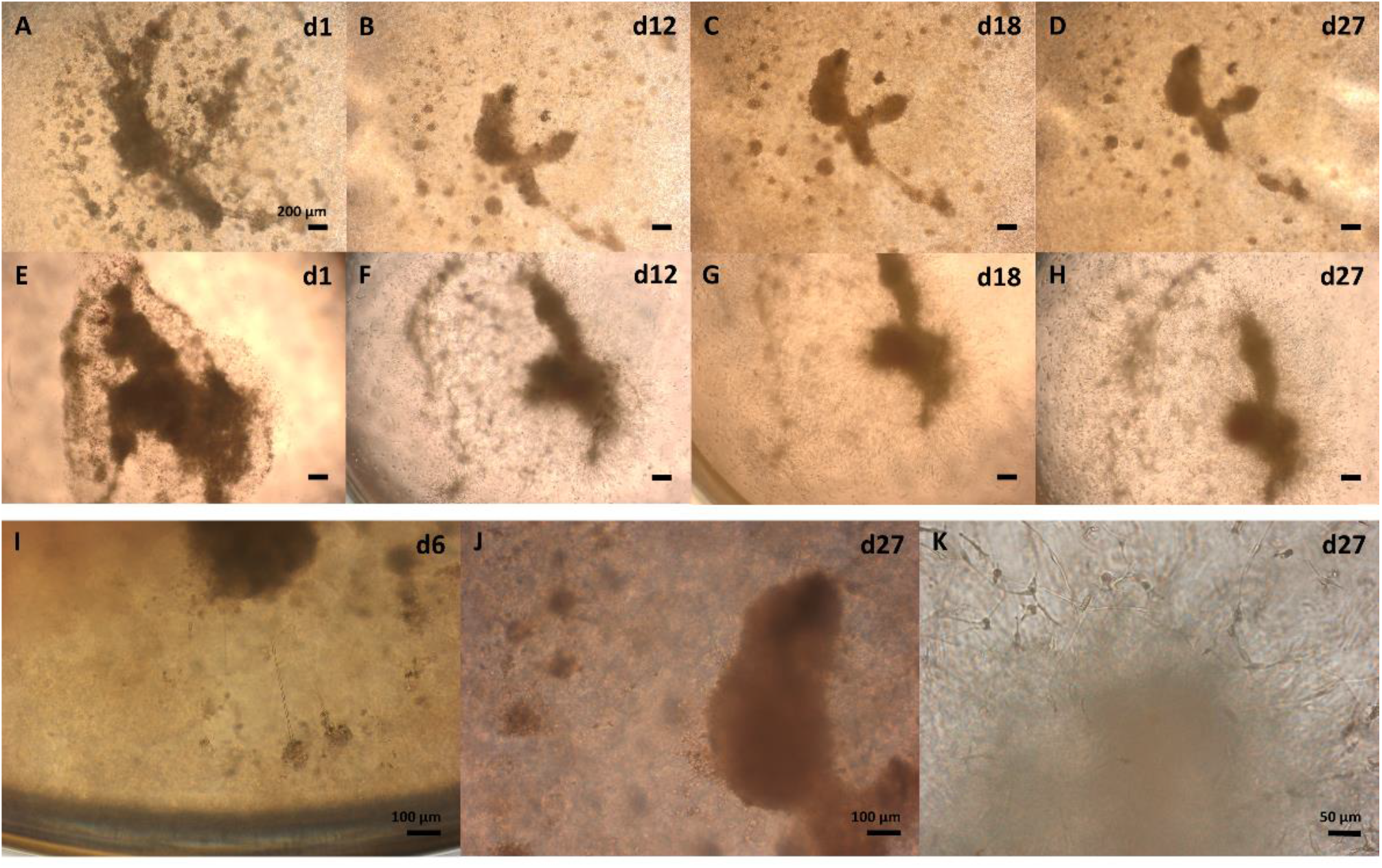
Morphology development of cultivated human glioblastoma multiforme (GBM) biopsy pieces in 0.7% nanofibrillar cellulose (NFC) and 40% Matrigel. Reductions in biopsy pieces’ size were common during NFC cultivation (A-D). New growth appeared on the earliest on day 6 in both scaffolds and consisted of protruding structures (I: NFC culture presented). Tissue pieces showed rapid spreading growth in surrounding Matrigel scaffolding, causing reductions to the original sample size (E-H). Observed novel growth consisted of protruding cells in both scaffold types (J: NFC; K: Matrigel). Figure scale bars 200 μm (A-H), 100 μm (I, J), 50 μm (K).

### Histological analysis of skin biopsies indicated active maintenance of tissue integrity when cultivated in NFC

HE analysis of ex vivo HS tissue piece revealed epidermal skin layers and dermal mesenchymal tissue (Fig 5A), and epidermal layers without mesenchymal tissue in NFC-cultivated samples (Fig 5D,C). The ex vivo HS tissue had strong immunoreactivity for mitotically active basal keratinocyte cytokeratins 5 and 6 (CK 5/6) in appropriate epidermal cell layers (Fig 5B), and displayed mediocre-level expression of intermediate filament type-III vimentin in epidermal spinous cell layer (Fig 5C). Cultivated HS tissue pieces sustained epidermal CK5/6 expression (Fig 5E,H) and displayed low vimentin expression in the spinous- (day 12,28) and cornified tissue layers (day 28) (Fig 5F,I).

**Fig 5.**
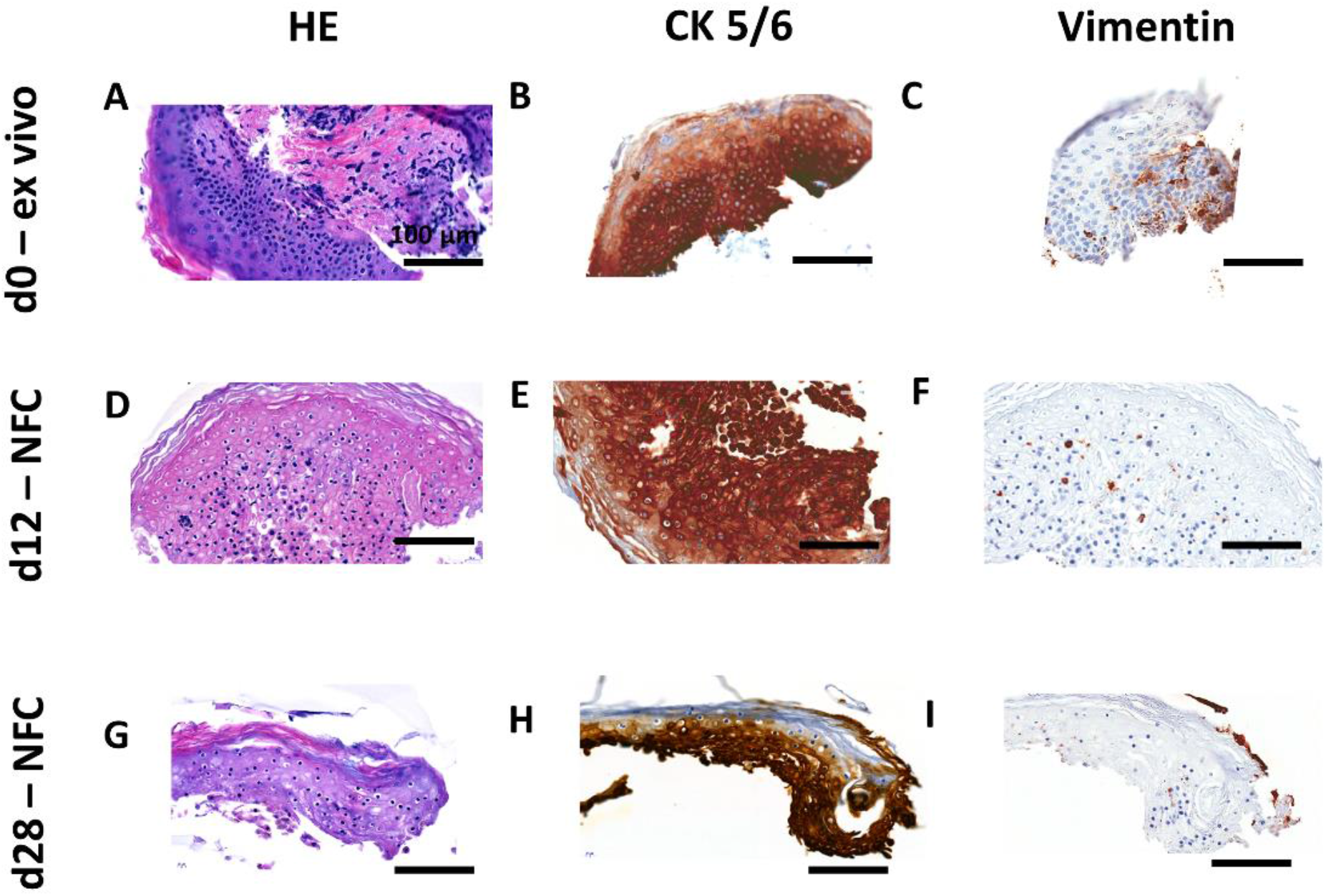
Histological hematoxylin-eosin (HE) tissue structure- and CK 5/6- and vimentin epitope analysis from healthy skin (HS) ex vivo- (day 0) and cultivated biopsy pieces (days 12, 28) in 0.7% nanofibrillar cellulose (NFC). HE analysis detected epidermal layers and dermal tissue in ex vivo HS sample (A), and epidermal tissue layers in day 12 (D), and day 28 samples (G). CK5/6 was detected in the epidermal cell layers of all analysed samples (day 0, 12, 28) (B, E, H). Vimentin stain was locating within the spinous layer of the ex vivo- (C) and day 12 and 28 samples (F, I), and additionally in the cornified layer of the day 28 sample (I). Figure scale bars 100 μm.

HE analysis of the ex vivo FRSK biopsy revealed epidermal layers and dermal tissue (Fig 6A). When comparing the ex vivo- and day 3 sample morphologies to further time points, it appeared that many biopsies were gaining spheroidical form during NFC cultivation. Matrigel-cultivated FRSK sample contained pink eosin protein formations encasing the darker hematoxylin-stained cell nuclei of the biopsy piece (Fig 6E). The ex vivo FRSK tissue had strong CK 5/6 staining in all epidermal layers with overlapping E-cadherin detection (Fig 6F,K), and mediocre-level vimentin throughout the biopsy piece (Fig 6P). During NFC cultivation, FRSK pieces displayed strong CK5/6 staining in apical cell layers (Fig 6G-I), low-level E-cadherin detection with some overlapping to CK 5/6 in NFC-2 and NFC-3 samples (Fig 6L-N), and from mediocre to strong vimentin expression in the interior parts of cultivated pieces (Fig 6Q-S). Due to samples’ size limitation for maximum number of analysis, positive vimentin had to be presented from another cultivated NFC-2 biological replicate (Fig 6R), that consisted of mesenchymal tissue and was lacking E-cadherin detection (Fig S10D). Matrigel-cultivated FRSK biopsy piece had nearly absent CK5/6 and E-cadherin detection (Fig 6J,O), and mediocre to strong vimentin expression in the interior parts of the biopsy piece (Fig 6T). Possible ruffling of sample specimens are indicated by black arrow point (Fig 6E,O,T).

**Fig 6.**
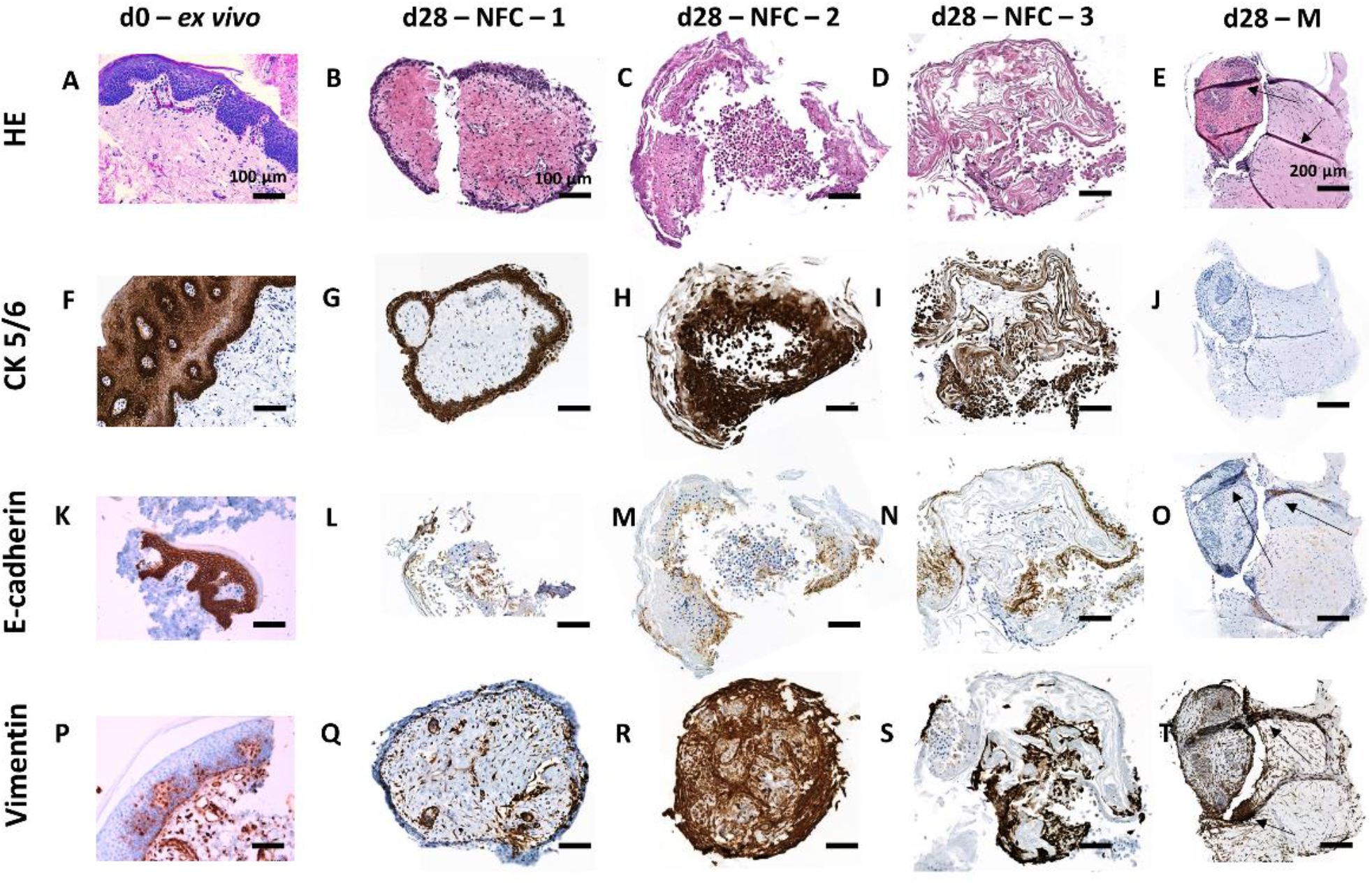
Histological hematoxylin-eosin (HE) tissue structure-, E-cadherin-, CK 5/6- and vimentin epitope analysis from healthy foreskin (FRSK) ex vivo- (day 0) and cultivated biopsy pieces (day 28) from three individual patients in 0.7% nanofibrillar cellulose (NFC) (as NFC-1,-2,-3) and 70% Matrigel (M). HE analysis revealed all epidermal layers and dermal tissue in the ex vivo FRSK biopsy piece (A). In NFC-cultivated day 28 samples, HE staining showed epidermal tissues in all samples and additional dermal tissue in NFC-1-, and possibly in NFC-3 replicate (B-D). Matrigel-cultivated biopsy piece was lacking clear tissue layers and consisted of darker hematoxylin-stained biopsy tissue surrounded by pink eosin-stained protein growth (E). Cytokeratin 5/6 was detected in the epidermal layers of the ex vivo sample (F) and in the outermost cell layers of NFC-cultivated biopsy pieces. (G-I). CK 5/6 was not detected in Matrigel-cultivated sample (J). Epithelial E-cadherin was detected locating to similar tissue layers with CK5/6 in ex vivo sample (K) and was positive in NFC-cultivated samples, with minor overlapping to CK 5/6 in NFC-2 and NFC-3 samples (L-N). E-cadherin was absent in Matrigel-cultivated sample (O). Vimentin was detected in the mesenchymal and occasional epidermal cells of the ex vivo biopsy sample (P), in the interior parts of NFC-cultivated biopsy pieces (Q-S), and was expressed throughout the whole biopsy piece in the Matrigel-cultivated sample (T). Ruffling of sample specimens are indicated by black arrow points (E, O, T). Figure scale bars 100 μm (d0-ex vivo column), 100 μm (NFC-cultivated sample columns) and 200 μm (Matrigel-cultivated sample column).

### Histological analysis of GBM cultures revealed sustained ex vivo protein expressions and novel expression patterns

HE analysis revealed spindle shaped, highly pleomorphic tumor cells in all analysed GBM samples (ex vivo, NFC, Matrigel) (Fig 7A,E,I). Antibody analysis revealed sustaining ex vivo protein expression profile in all culture conditions (NFC, Matrigel) until day 28. Immunoreactivity to oligodendrocyte transcription factor 2 (Olig2) had evenly distributed nuclear stain from ex vivo to day 28 in vitro (Fig 7B,F,J). Similarly, calcium ion binding protein B (S100) detection remained stable from ex vivo to day 28 in vitro (Fig 7C,G,K). Neuroectodermal stem cell - intermediate filament VI (Nestin) appeared during cultivation and was detected in cultivated day 28 samples (Fig 7D,H,L).

**Fig 7.**
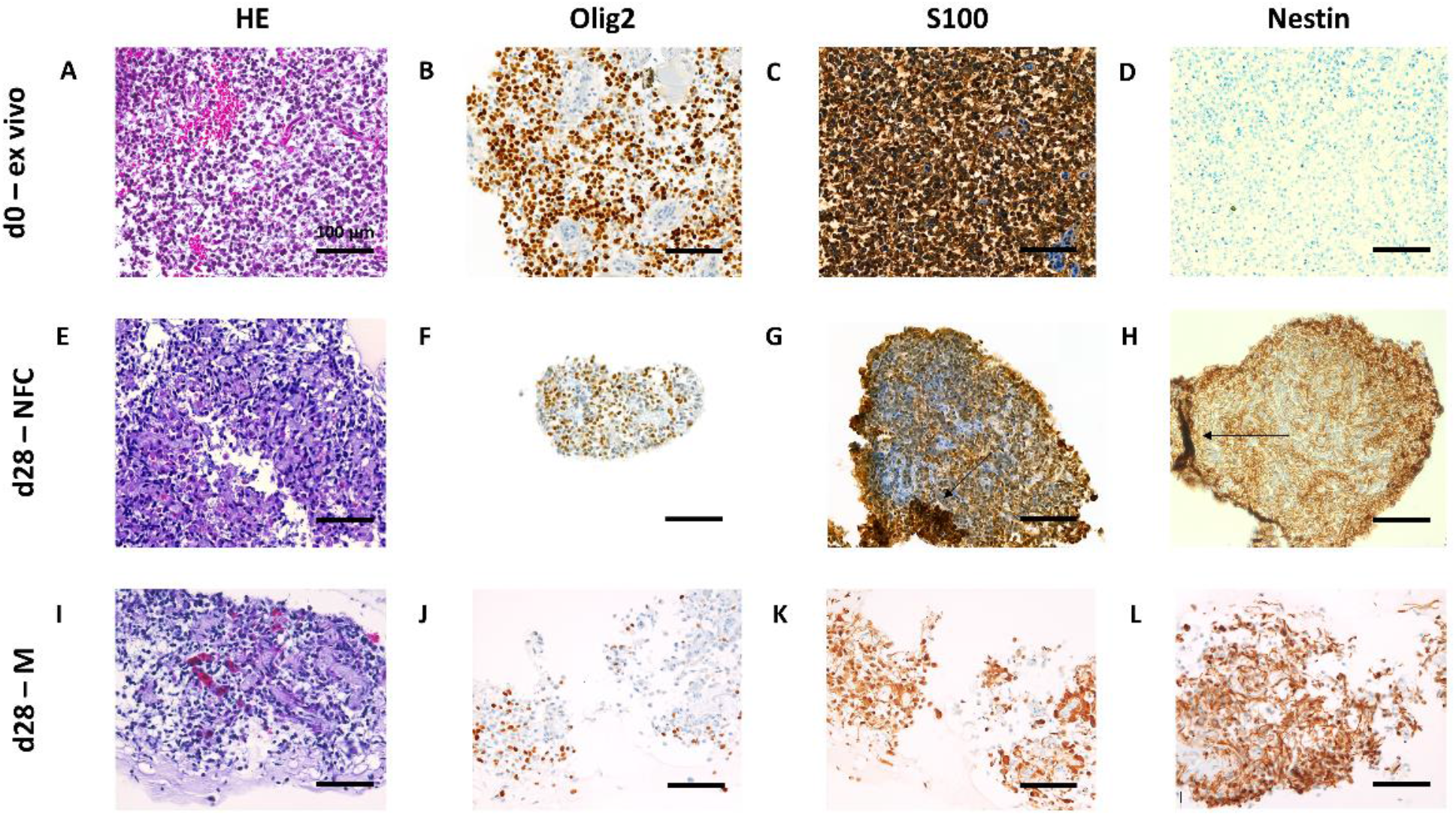
Histological hematoxylin-eosin (HE) tissue structure- and Olig2, S100 and Nestin epitope analysis from glioblastoma (GBM) ex vivo- (day 0) and cultivated biopsy pieces (day 28) in 0.7% nanofibrillar cellulose (NFC) and 40% Matrigel (M). HE analysis revealed pleiomorphic tumor cell structure in ex vivo- and cultivated day 28 GBM pieces (A, E, I). Olig2 showed equally distributed detection stain in all samples from ex vivo- to NFC- and Matrigel cultivated day 28 GBM samples (B, F, J). S100 was similarly detectable already in the ex vivo biopsy piece and remained stable during cultivation in both culture conditions (C, G, K). Nestin was absent in ex vivo sample (D) and appeared during cultivation in both culture conditions (H, L). Ruffling of sample specimen is indicated by black arrow points (G, H). Figure scale bars 100 μm.

Also transmembrane ligand receptor NOTCH1 appeared during cultivation and had peripheral location in NFC-cultivated day 6, 12 and 28 samples (Fig 8A,D). NOTCH1 was positive also in Matrigel-cultivated samples and showed possible accentuation of staining in day 12 sample edges, but not in day 28 sample (Fig 8G). Immunoreactivity to microtubule-associated protein 2 (MAP2) had partly perinuclear location in ex vivo and day 28 samples, and showed peripheral staining in NFC-cultivated sample piece (Fig 8B,E,H). Astrocyte intermediate filament-specific glial fibrillary acidic protein (GFAP) had strong detection in ex vivo- and day 28 samples (Fig 8C,F,I), and showed peripheral staining in NFC-cultivated sample piece edges on day 28- and possibly in day 12 sample, but not in day 6 sample (Fig 8F), or any Matrigel-cultivated samples (Fig 8I). Ruffling of sample specimens are indicated by black arrow point (Fig 8G,H).

**Fig 8.**
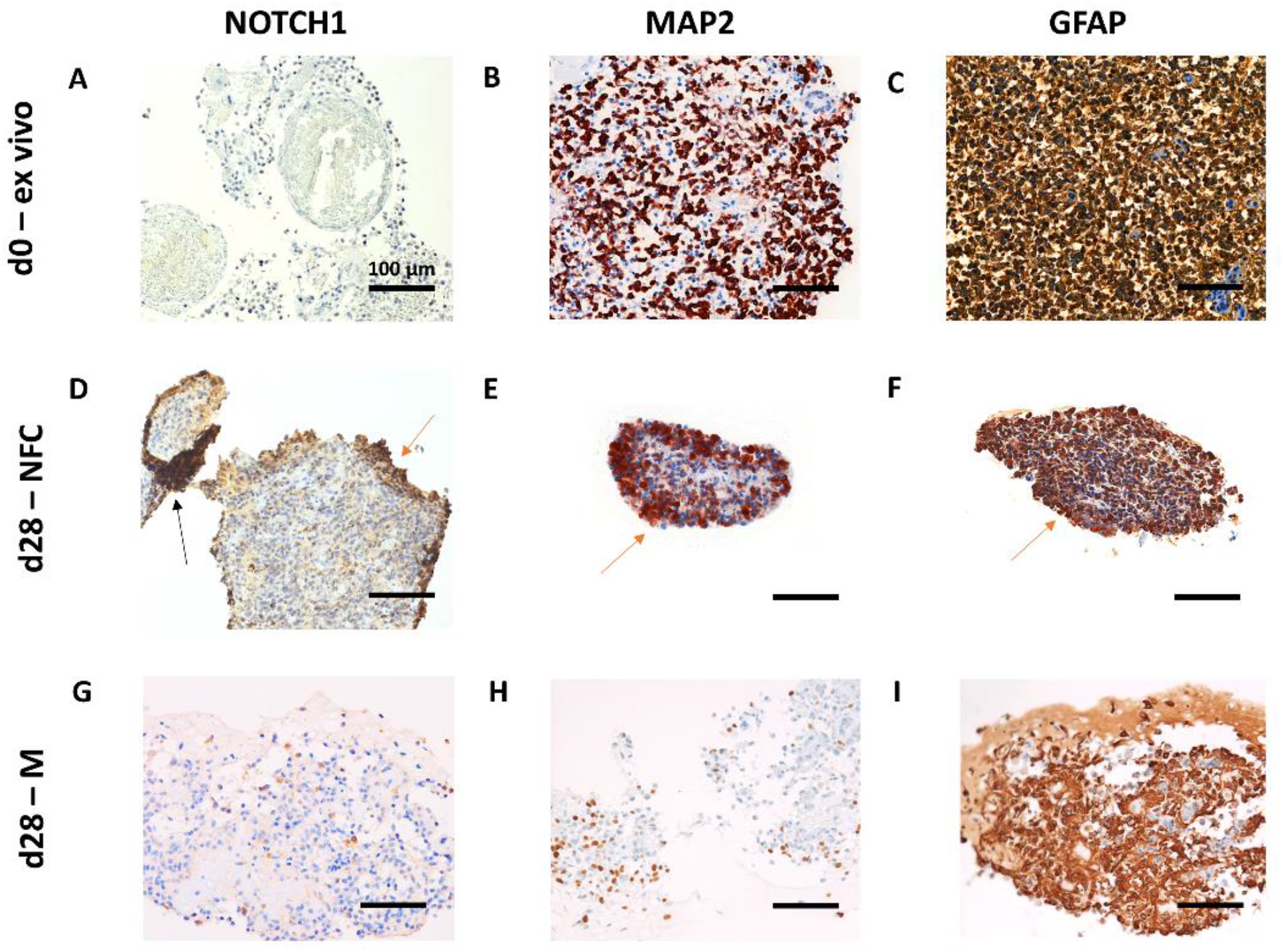
Histological NOTCH1, MAP2 and GFAP epitope analysis from glioblastoma (GBM) ex vivo- (day 0) and cultivated biopsy pieces (day 28) in 0.7% nanofibrillar cellulose (NFC) and 40% Matrigel (M). The expression of NOTCH1 was absent in the ex vivo sample (A) and appeared during cultivation in both culture conditions with peripheral staining in NFC-cultivated sample (D), and barely detectable expression in Matrigel– cultivated sample (G). MAP2 had stable detection in ex vivo sample (B), peripherally accentuated staining in day 28 NFC sample (E), and weaker detection in Matrigel-cultivated sample (H). GFAP expression was detected in the ex vivo sample piece (C) and remained stable during cultivation in both NFC and Matrigel conditions with minor peripheral accentuation in NFC –cultivated sample (F,I). Accentuated staining patterns are indicated by orange arrow points (D, E, F). Ruffling of sample specimen is indicated by black arrow point (D). Figure scale bars 100 μm (A-I).

## Discussion

In this study, we developed technically efficient methodology to sustain intact and heterogeneous human tissues long-term in vitro using NFC scaffolding. Parallel Matrigel tissue cultures were done to provide a commonly used comparative 3D scaffold that is well known for supporting any 3D growth.

Structural HE analysis of the skin tissues revealed differing behaviors between HS and FRSK biopsies in vitro. Where HS biopsies sustained relatively static tissue structure during cultivation, FRSK culture repeats indicated re-organizations in tissue structure. In order to verify biopsies’ morphology development as cell movement as well as to characterize present cell types, both skin tissue types were evaluated for their distributions of vimentin. The ex vivo- and NFC-cultivated HS tissues as well as ex vivo FRSK tissue demonstrated randomly distributed and partially epidermal low-level vimentin stain, which can be considered as an indicator of epidermal melanocytes as well as activated keratinocytes during wound healing [23,24]. During NFC- and Matrigel cultivation, HS biopsies showed low vimentin detection, and FRSK biopsies mediocre-to-strong vimentin detection. The detected clear vimentin stain in FRSK cultures, but not in HS cultures, could therefore support the visual observations of ongoing morphology alterations in cultivated FRSK pieces as vimentin enables cell motility [25,26]. Differences in HS and FRSK vimentin expressions may reflect differing characteristics of the two tissue-types, or originate from heterogeneous structural variation in sample compositions; e.g. the quantities of mesenchymal tissue and fibroblasts required for biopsy’s representative “wound” closure [26]. In addition to vimentin expression, HS and FRSK samples were evaluated for their expression of CK 5/6 that is typical for basal keratinocytes maintaining structural integrity of the epidermis [27,28]. FRSK samples were also evaluated for the expression of E-cadherin that indicates stable adherens junctions of immobile keratinocytes [29,30]. The extensive epidermal CK 5/6 staining in all HS samples indicated normal tissue structure and active maintenance of tissue’s structural integrity. Strong CK 5/6 expression was also detected in ex vivo and NFC-cultivated FRSK samples, where it was correctly localizing in the epidermal layers of the ex vivo FRSK biopsy piece, and was found encasing the sur-faces of spheroidically organizing biopsy pieces during cultivation. Positive E-cadherin was accordingly detected overlapping with CK 5/6 expression in the ex vivo FRSK sample, and similarly on sample surfaces in the NFC-cultivated tissue pieces. The expressional overlapping of E-cadherin and CK 5/6 was less evident in cultivated FRSK samples, which might be, similarly to vimentin expression, due to the differences in biopsies’ tissue compositions, or the natural lag in keratinocytes’ E-cadherin expression during wound healing [31,29]. These findings suggest the survival of HS and FRSK tissues’ natural characteristics during NFC cultivation, and in the case of FRSK tissue, indicate an ability to re-organize into spatially correct cell layers in vitro [30]. Judging by the lacking CK5/6- and E-cadherin- and positive vimentin expressions, Matrigel condition promoted migratory behavior in biopsy’s cells. The observed eosin formations around Matrigel-cultivated biopsy pieces might also indicate wound healing-specific fibroblast activity in the form of collagen production [26]. Taken together, skin biopsy cultures could provide biologically relevant material for the modelling of wound healing and pathological conditions [15, 32], and for the experimental introduction of various pro-inflammatory or therapeutic factors, carcinogens, pathogens or metastasizing cancer cells [14].

As the neoplastic biopsy model of this study, GBM patient tissue was utilized. HE analysis revealed that typical histopathological GBM tumor tissue structure was sustained during NFC and Matrigel cultivation. The ex vivo- and NFC- and Matrigel-cultivated glioblastoma biopsy pieces stained positive for the characterized Olig2, S100, Nestin, NOTCH1, MAP2 and GFAP biomarkers, indicating successfully preserved brain tumor tissue in vitro. According to the acquired profiling, the biopsy pieces contained neoplastic glial cells or mobile immature glial cells expressing MAP2, typical for structural stabilization during neuronal development [33]; mature astrocytes or glial astrocytes (GFAP) [34,35], expressing S100 concentration-dependent proliferation, neurite outgrowth, or inflammatory- and neoplasia -related activity [36,35,37], and stem cells, indicated by Olig2 that has been reported to sustain brain tumor stem cell populations [38]. The expressions of general upstream-regulator NOTCH1 and Nestin are also common in brain neoplasia, where NOTCH1 maintains and drives the proliferation of cancer stem cells [39], and Nestin marks differentiating neural progenitor cells [40]. In addition, NFC-cultivated glioblastoma tissues showed possible signs of reactive gliosis, observed as peripheral staining patterns of NOTCH1, MAP2 and GFAP in the biopsy pieces [41,42,43]. In non-neoplastic tissues, gliosis is triggered by tissue trauma factors, such as growth factors released from damaged tissue, factors presented by direct blood contact, and descending oxygen and nutrient levels after cutting off blood circulation [44]. However, during cancer progression, many stem cell-specific-, angiogenic- and growth-related genes are expressed out of their developmental context and are enabling tumor progression [45,46], from which neoplastic gliosis is a commonly recognized example of [47,48]. Matrigel cultivated pieces demonstrated equally strong expressions of S100, Nestin and GFAP biomarkers, had significantly weaker expressions of Olig2, NOTCH1 and MAP2, and showed no indications of gliosis. This suggests less variation in surviving cell populations in vitro, and alterations in the original tissue expression profile. Comparing to cur-rent brain tissue research, the induction of GFAP expression in vitro generally requires additional nitrogen oxide or acidic medium compounds [49,50]. As all GBM cultures demonstrated stable GFAP expression in this study, the established biopsy culture models might represent slightly acidic or hypoxic long-term growth environment, reminiscent of tumorigenic or inflamed brain tissues [49,50,41,51]. In addition to possible hypoxia and neoplastic gliosis, the observed peripheral staining patterns of NOTCH1, GFAP and MAP2 may also originate from physiological responses to ischemic tissue areas caused by sample dissection [41,42,44,52,53]. The lacking expressions of NOTCH1 and Nestin in the ex vivo GBM biopsy could be due to natural lag in post-injury cell responses. For example, NOTCH1 has been detectable after 12-24h post-trauma in the rat models [53]. Due to the presented natural tissue responses in vitro, NFC-brain biopsy cultures may offer biologically relevant material for the modelling of gliosis [44,54], various pathological conditions [44,45,51] or introduction of therapeutic factors [55,52]. In addition, since maturation of stem cell –derived neural cell types in vitro is time-consuming and technically very demanding even with one successful cell type [56,57], ex vivo brain cultures could provide efficient alternative for stem cell -derived models.

The NFC-, and to an extent, also Matrigel culture environment, appeared consistent in preserving the histopathological traits of patient biopsies. The observed cell outgrowth and developments in tissue morphology also indicated culture viability. This conclusion could be supported by the steady vimentin stain in FRSK culture repeats, the detected clear nuclei and tissue structure in all HE analysis, and by the continuous S100 detection in the glioblastoma cultures. Although the value of S100 expression, typical for proliferating tissues, has been found contradictory as an excluding biomarker for necrosis [58,59], vimentin has been generally recognized to be degraded in apoptotic cells [60,61], and round eosin staining patterns common for necrotic tissues, were absent in analysed samples. When comparing the two culture scaffolds, 3D Matrigel appeared to promote cell migration in the expense of tissue integrity in FRSK and GBM cultures, which was also marked by the losses of tissue heterogeneity and typical gliosis processes in GBM histology. Indeed, although considered as the golden standard in many 3D culture applications for decades, the use of Matrigel has been disputed due to included growth-guiding- and cancerous growth factors in experimental settings [62,63,64], as well as the fact that laminin-111 is crucial mostly during tissue development and expressed only in few tissue epithelia in adults [65]. We consider the combination of biopsy tissue and NFC hydrogel a very promising 3D model, since NFC matrix ensures that no unknown growth-affecting factors arise from the scaffold material and that the detected animal proteins originate exclusively from the cultured cells or known medium supplements. Long-term NFC cultivation of human samples could therefore be used for various research questions, be it modelling different disease models, drug screening or even personalized medicine.

## Conclusions

We report here the development of novel in vitro tissue cultivation methodology utilizing nanofibrillar cellulose (NFC; GrowDex) hydrogel; including the 3D culture assembly, optimized maintenance protocol and sample preservation methodology for NFC-cultivated biopsy pieces. The results of histopathological profiling from healthy skin-, healthy fore-skin- and neoplastic glioblastoma multiforme biopsies indicated that biopsy pieces survive in long-term NFC cultivation and remain capable of sustaining their histological tissue identities. In addition to tumor modelling, we hypothesize this novel tissue culture model applicable to the modelling of regenerative and pathological processes that share many molecular features and regulatory mechanisms with cancers. Further studies for the characterization of surviving cell-types long-term in vitro as well as prevailing oxygen- and pH conditions in long-term NFC culture niche would further promote the understanding of the possibilities and limitations of this novel tissue culture system.

## Acknowledgments

This work was supported by UPM-Kymmene, Finland. We thank the many researchers and staff in the Core Facility Alternative Biomodels and Preclinical Imaging and Diagnostic and Research Institute of Pathology at the Medical University of Graz for their extra time and effort given during the methodology development.

